# Laboratory culture of the California Sea Firefly *Vargula tsujii* (Ostracoda: Cypridinidae): Developing a model system for the evolution of marine bioluminescence

**DOI:** 10.1101/708065

**Authors:** Jessica A. Goodheart, Geetanjali Minsky, Mira N. Brynjegard-Bialik, Michael S. Drummond, J. David Munoz, Timothy R. Fallon, Darrin T. Schultz, Jing-Ke Weng, Elizabeth Torres, Todd H. Oakley

**Affiliations:** Department of Ecology, Evolution, and Marine Biology, University of California, Santa Barbara, Santa Barbara, CA 93106; Department of Biological Sciences, California State University, Los Angeles, CA 90032-8201, USA; Whitehead Institute for Biomedical Research, Cambridge, MA, 02142, USA; Department of Biology, Massachusetts Institute of Technology, Cambridge, MA 02142, USA; Monterey Bay Aquarium Research Institute, Moss Landing, CA 95060, USA; Department of Biomolecular Engineering and Bioinformatics, University of California, Santa Cruz, Santa Cruz, CA 96060, USA

**Keywords:** ostracod, cypridinid, animal culture, cypridinid luciferin, embryogenesis, Crustacea

## Abstract

Bioluminescence, or the production of light by living organisms via chemical reaction, is widespread across Metazoa. Culture of bioluminescent organisms from diverse taxonomic groups is important for determining the biosynthetic pathways of bioluminescent substrates, which may lead to new tools for biotechnology and biomedicine. Some bioluminescent groups may be cultured, including some cnidarians, ctenophores, and brittle stars, but those use luminescent substrates (luciferins) obtained from their diets, and therefore are not informative for determination of the biosynthethic pathways of the luciferins. Other groups, including terrestrial fireflies, do synthesize their own luciferin, but culturing them is difficult, and the biosynthetic pathway for firefly luciferin remains unclear. An additional independent origin of endogenous bioluminescence is found within ostracods from the family Cypridinidae, which use their luminescence for defense and, in Caribbean species, for courtship displays. Here, we report the first complete life cycle of a luminous ostracod (*Vargula tsujii* Kornicker & Baker, 1977, the California Sea Firefly) in the laboratory. We also describe the late-stage embryogenesis of *Vargula tsujii* and discuss the size classes of instar development. We find embryogenesis in *V. tsujii* ranges from 25-38 days, and this species appears to have five instar stages, consistent with ontogeny in other cypridinid lineages. We estimate a complete life cycle at 3-4 months. We also present the first complete mitochondrial genome for *Vargula tsujii*. Bringing a luminous ostracod into laboratory culture sets the stage for many potential avenues of study, including learning the biosynthetic pathway of cypridinid luciferin and genomic manipulation of an autogenic bioluminescent system.

## Introduction

The production of bioluminescence in animals is an important biological phenomenon that is particularly prevalent in marine environments ^1^. To produce light, animals combine an enzyme (usually a luciferase or photoprotein) and a substrate (luciferin) with oxygen, and sometimes additional components such as ATP or calcium ^2–4^. Although the enzymes that produce light are well-studied ^5–13^, and we know how the substrate is produced in bacteria ^14^ and fungi ^15^, we do not yet know how luciferin is synthesized in any animal system. A comprehensive understanding of this pathway could be transformative for optogenetics in transgenic organisms, where the ability to produce bioluminescence in situ as a biosensor or light source would be invaluable. Some laboratory-reared luminescent animals already exist (Table 1), but may not be suitable for studying luciferin biosynthesis. In particular, ctenophores, cnidarians, and brittle stars use coelenterazine obtained from their diet as a substrate ^1^. Fireflies do use an endogenous luciferin specific to Coleoptera (beetles) ^16^, but they often have longer life cycles than other taxa, slowing research progress ^17^. Although there are other independent origins of autogenic bioluminescence, which we define as that produced via luciferase and luciferin that are synthesized via the genome of the light producing animal, no laboratory cultures of these animals currently exist. This makes additional studies focused on the comparative evolution of this ability extremely challenging.

**Table 1.**
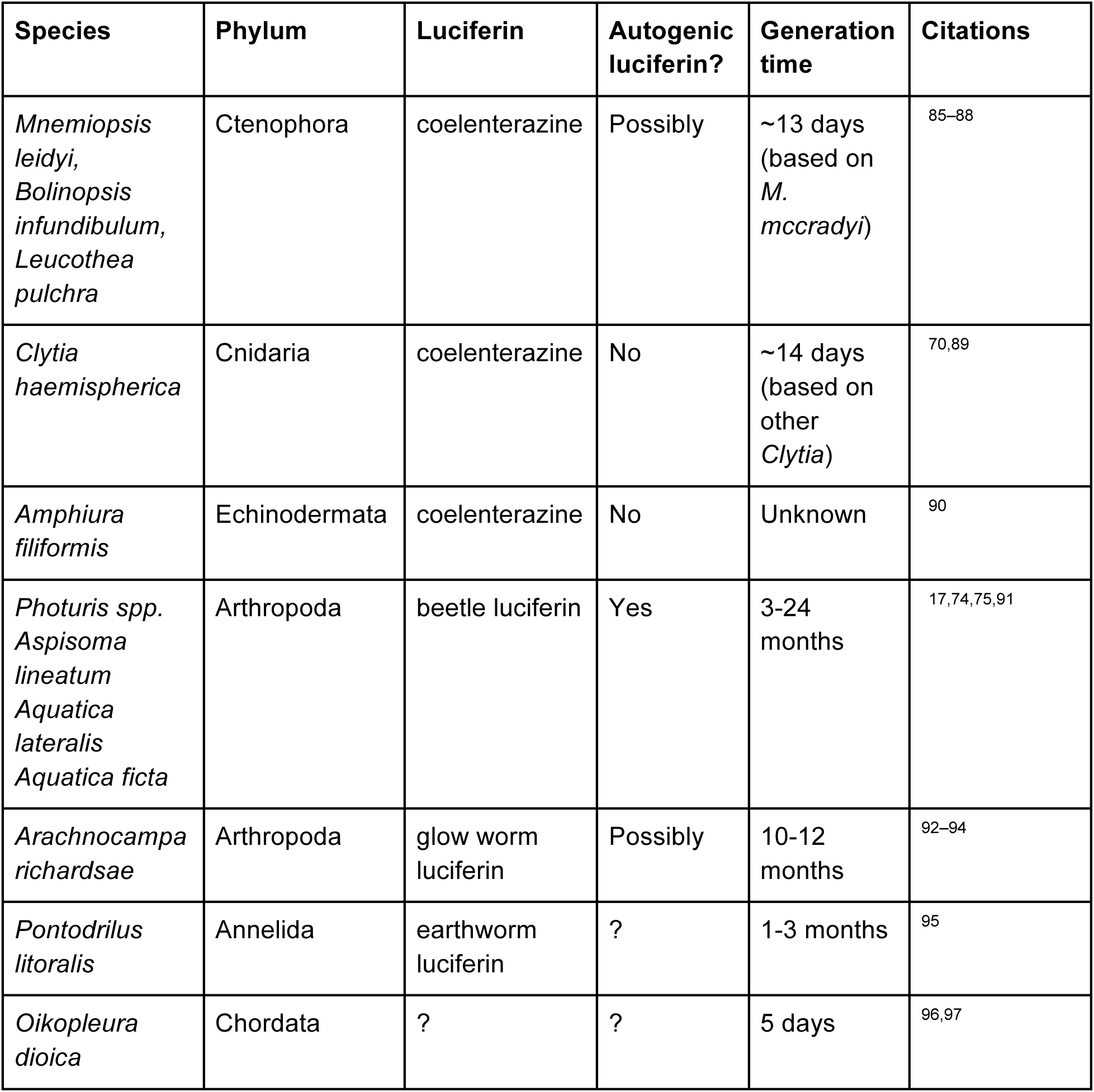
Species of bioluminescent metazoans for which cultures are known to exist, including information on the luciferin used for luminescence, estimated generation times, and whether the luciferin used is synthesized via the genome of the luminescent organism itself (autogenic).

| Species | Phylum | Luciferin | Autogenic luciferin? | Generation time | Citations |
| --- | --- | --- | --- | --- | --- |
| *Mnemiopsis leidyi*,<br>*Bolinopsis infundibulum*,<br>*Leucothea pulchra* | Ctenophora | coelenterazine | Possibly | ~13 days<br>(based on *M. mccradyi*) | 85–88 |
| *Clytia haemispherica* | Cnidaria | coelenterazine | No | ~14 days<br>(based on other *Clytia*) | 70,89 |
| *Amphiura filiformis* | Echinodermata | coelenterazine | No | Unknown | 90 |
| *Photuris* spp.<br>*Aspisoma lineatum*<br>*Aquatica lateralis*<br>*Aquatica ficta* | Arthropoda | beetle luciferin | Yes | 3-24 months | 17,74,75,91 |
| *Arachnocampa richardsae* | Arthropoda | glow worm luciferin | Possibly | 10-12 months | 92–94 |
| *Pontodrilus litoralis* | Annelida | earthworm luciferin | ? | 1-3 months | 95 |
| *Oikopleura dioica* | Chordata | ? | ? | 5 days | 96,97 |

**Table 2.**
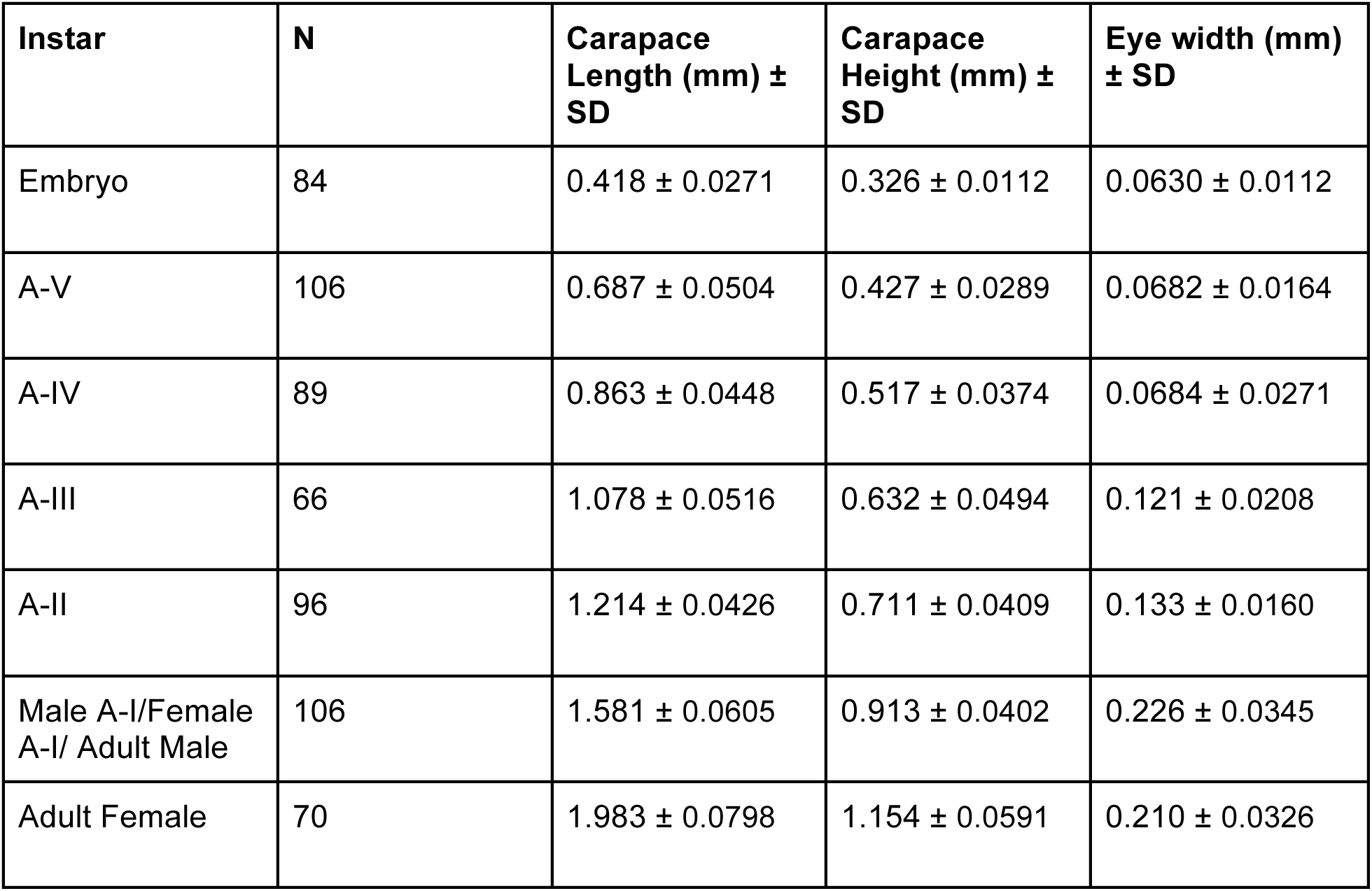
Measurements for each cluster (and inferred instar) of *Vargula tsujii* ± SD.

Ostracods from the family Cypridinidae independently evolved bioluminescence, and use a luciferin that is endogenous to ostracods and chemically different from that of other bioluminescent organisms ^2, 18, 19^. Of the metazoan taxa already cultured, autogenic luminescence is only known to occur within fireflies ^20, 21^. Due to the independent origin of ostracod bioluminescence in the family Cypridinidae, and evidence for the autogenic nature of the necessary luciferin and luciferase, ostracods represent an importantl lineage for understanding how bioluminescence evolved. Establishing laboratory cultures of luminescent ostracods within Cypridinidae will provide opportunities for new investigations into the biosynthetic pathway of luciferins in Metazoa, as well as comparative studies of the biochemistry, physiology, genetics, and function of endogenous bioluminescence.

Cypridinidae is estimated to include over 300 ostracod species ^22^, some of which use bioluminescence for defense, while others possess the ability to create bioluminescent courtship displays in addition to defensive secretions ^23^. Lineages with sexually dimorphic bioluminescent displays, such as those in Cypridinidae, have more species than their non-displaying sister lineages, indicating that origins of bioluminescent courtship may increase diversification rates ^24^. As such, cypridinid ostracods represent an excellent system for understanding how behavioral diversification can lead to increases in species accumulation. Furthermore, cypridinid ostracods diversified while generating bioluminescence with a consistent biochemical basis across the clade ^25^, meaning that they use the same luciferin and homologous enzymes to make light. The conserved nature of the luminescent chemical reaction provides a system within which bioluminescence can produce rapid changes in phenotypic diversity by modifying amino acid sequences of the enzyme and how the enzyme is used ^26^.

Attempts to culture luminescent and non-luminescent species of Cypridinidae have led to varying success, including *Skogsbergia lerneri* (non-luminous species with no mating in culture) ^27^, *Photeros annecohenae* (luminous species without the complete life cycle in culture) ^28^, and *Vargula hilgendorfii* (luminous species with no mating in culture) ^29^. Researchers attempting to culture *Vargula hilgendorfii* tried multiple alternative aquarium set-ups and tested sensitivity to nutrients commonly measured in aquaria, but were still unable to induce mating in the lab ^29^. An unnamed species of luminescent ostracod is claimed to have been cultured at the CTAQUA Foundation (Aquaculture and Technological Centre) in Spain for molecular gastronomy ^30^, but no information on species, culture longevity, or whether mating occurred is available on this enigmatic and uncertain culture. Other non-cypridinid ostracods have been reared in the laboratory as well, including the myodocopids *Euphilomedes carcharodonta* ^31^ and *Euphilomedes nipponica* ^32^ and the podocopids *Xestoleberis hanaii* and *Neonesidea oligodentata* ^33^ (only records of copulation in the laboratory for any ostracods) ^34^. Many podocopids grow quite easily in aquaria, so a number of other species have been cultured for descriptions of, or experiments related to, post-embryonic development, including *Chlamydotheca incisa* and *Strandesia bicuspis* ^35^*, Heterocypris salina* ^36–38^*, Eucypris virens* ^37^, and *Candona rawsoni* ^38^.

In this paper, we present evidence that we are the first to rear a luminous ostracod (*Vargula tsujii* Kornicker & Baker, 1977) through its complete life cycle in the laboratory. *Vargula tsujii* is the only bioluminescent cypridinid ostracod described from the Pacific coast of North America. Commonly found in southern California, the reported range of *V. tsujii* is from Baja California to Monterey Bay ^39^. This species is demersal (moves between the benthic and planktonic environments) and is primarily active in or near the kelp forest at night ^40^. Similarly to some other cypridinid ostracods, *Vargula tsujii* is a scavenger, and is presumed to feed on dead and decaying fish and invertebrates on the benthos since they can be collected via baited traps. In addition, some authors observed *V. tsujii* attacking the soft tissues of live fishes caged nearshore ^41^. *Vargula tsujii* is an important prey item of the plainfin midshipman fish (*Porichthys notatus*), as it is the source of luciferin for the bioluminescence seen in the fish ^42^.

In addition to evidence for the completion of the life cycle of *Vargula tsujii* in the laboratory, we provide a successful methodology for culturing populations of *V. tsujii*, descriptions and analyses of late stage embryogenesis and instar development, a reconstruction of its mitochondrial genome, and a brief description of the distribution of molecular diversity of *V. tsujii* across coastal southern California.

## Methods

### Specimen collection

We collected *Vargula tsujii* from three primary locations at 0-10 m depth: (1) Fisherman’s Cove near Two Harbors on Catalina Island, CA, USA (33°26’42.5″N 118°29’04.2″W) directly adjacent to the Wrigley Marine Science Center, (2) Cabrillo Marina, Los Angeles, CA, USA (33°43’13.5″N 118°16’47.4″W), and (3) Shelter Island public boat ramp, San Diego, CA, USA (32°42’56.3″N 117°13’22.9″W) (Figure 1; Table S1). We also provide locations in Figure 1 and Table S1 where we set traps but did not collect *V. tsujii*. The specimens used for culture and description of the life cycle of *V. tsujii* were exclusively from Fisherman’s Cove, Catalina Island, and we collected additional specimens from the Cabrillo Marina and Shelter Island public boat ramp to discern the genetic diversity of *V. tsujii* and test for the presence of any cryptic species. For specimen collection we used Morin pipe traps: cylindrical polyvinyl chloride (PVC) tubes with mesh funnels to prevent the ostracods from escaping once they entered ^43^. We baited traps with dead fish or fresh chicken liver (roughly 1-2 cm^3^). We set traps in 1-10 meters of water at sunset and collected them 1.5 to 2 hours after last light to allow time for the ostracods to accumulate.

**Figure 1.**
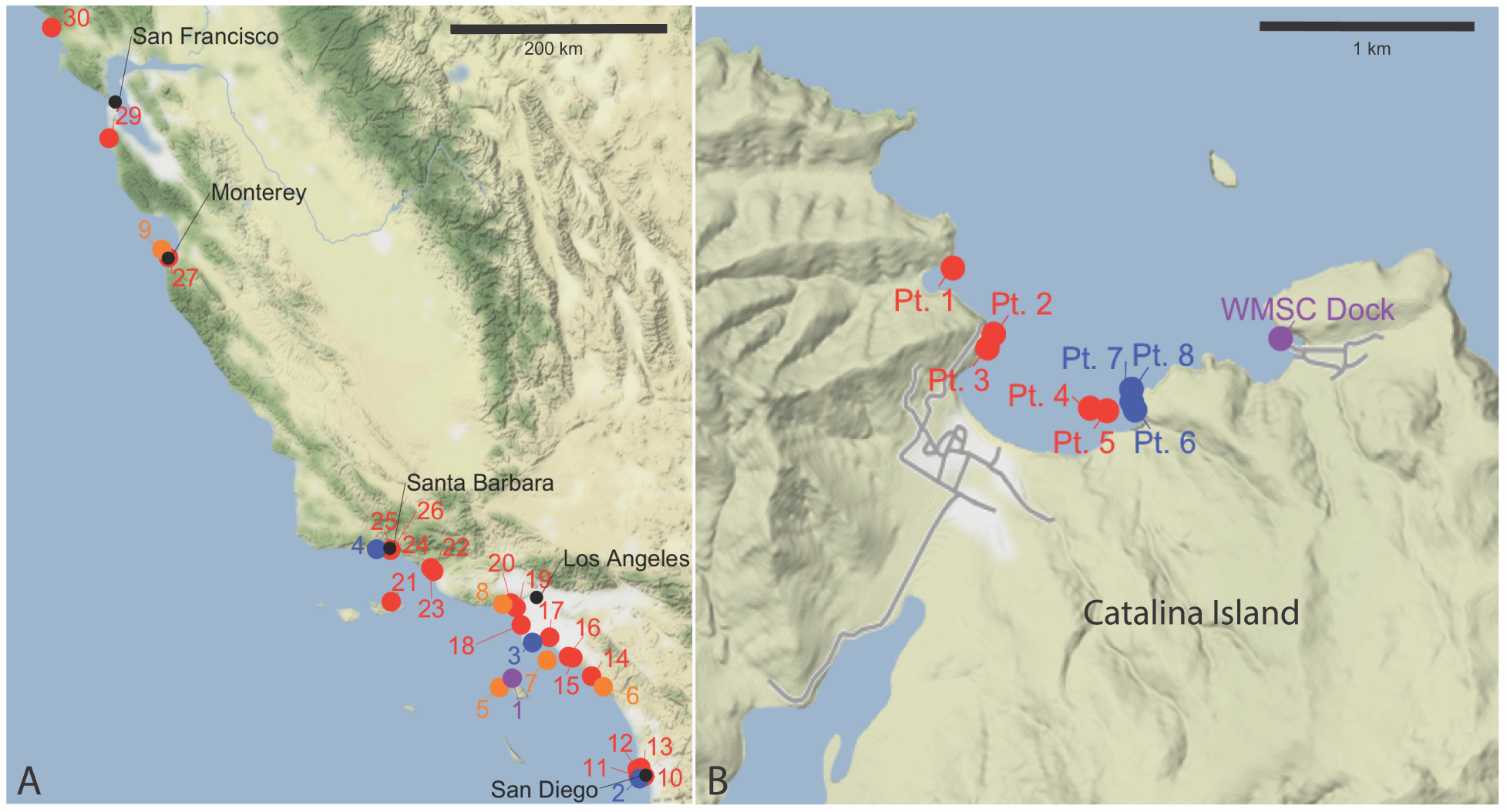
Map of localities where collections (and attempted collections of *Vargula tsujii* were made). Black points represent geographic annotations for reference, purple points refer to the location where ostracods were collected that were ultimately cultured, blue where animals were found or collected, orange where Kornicker originally collected animals, and red where no *V. tsujii* were found. (A) Map of the California (USA) coastline where V. tsujii collections were attempted; (B) Map of the region near Two Harbors (Santa Catalina Island, California, USA) where V. tsujii collections were attempted.

### Population genetic analyses

We obtained *Vargula tsujii* sequence data for 16S rRNA (16S) and cytochrome oxidase 1 (COI) from a previously published thesis ^44^. We then aligned and visually inspected sequence chromatograms with Sequencher version 3.1 (Gene Codes Corp) (Table S1). We constructed haplotype networks for the two genes, COI (Number of individuals: Catalina, 18; San Diego, 11; San Pedro, 12) and 16S (Number of individuals: Catalina, 19; San Diego, 20; San Pedro, 21) with the package pegas ^45^ in the R environment (version 3.4.2) ^46^. To assess whether sequences within populations cluster together, we analyzed each gene via Discriminant Analysis of Principal Components (DAPC) using the adegenet package in R ^47^ and compared models with one, two, and three clusters using the Bayesian Information Criterion (BIC) ^48^.

### Description of aquarium system and feeding

We cultured *Vargula tsujii* at the University of California, Santa Barbara in aquariums designed for water to flow through sand and out the bottom. This allowed continuous water flow to keep water fresh, and not suck ostracods into a filter. Instead, ostracods remain in sand where they spend the daylight hours. We kept animals under a 12:12 light cycle with an indirect blue LED light.

We first modified a five gallon glass tank by drilling a hole in the bottom, which was then covered with the bottom of an undergravel filter and a 3 cm layer of commercial aragonite sand with grain sizes 1-2 mm for aquariums (CaribSea brand). To maintain continuous water flow, we used plastic tubing (diameter = 1 cm) connected directly to an overflowing reservoir tank, itself fed by UCSB’s seawater system that pumps directly from the Pacific. The tube from the reservoir created a siphon system for moving water into the main tank (Figure 2). The siphon provided continuous water flow when needed, and prevented overflow of the aquarium. In addition, to prevent a complete drain of the water if the siphon flow broke, which is common with larger siphons, we attached an outflow pipe to the bottom of the aquarium that drained halfway up the side of the tank (Figure 2).

**Figure 2.**
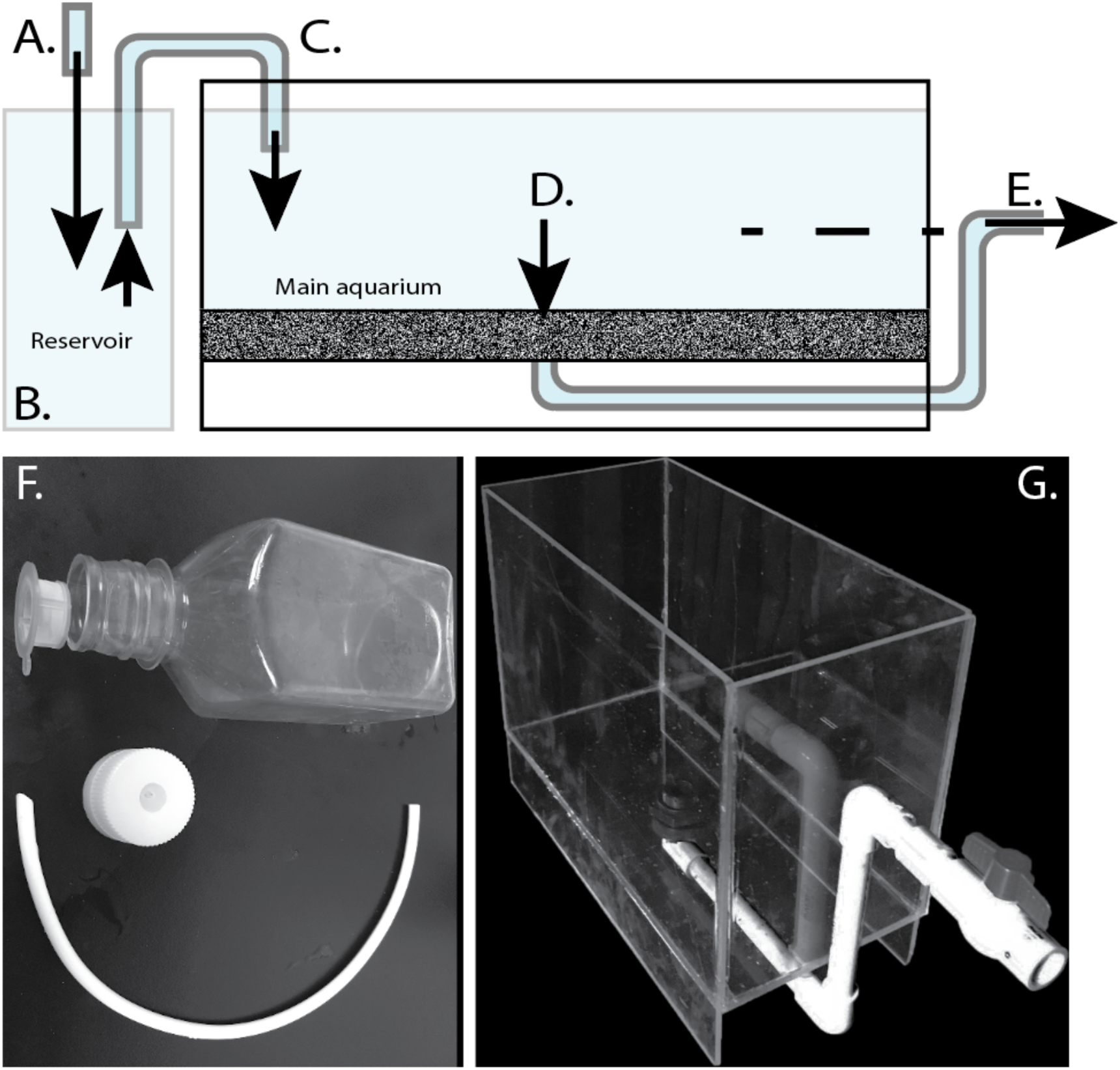
Diagrammatic representation of aquarium setup for Vargula tsujii. (A) Source water flows into (B) a reservoir tank, which overflows to keep the reservoir full and water level constant. (C) We use a siphon to carry water from a reservoir tank to main aquarium where ostracods live. Water level of the main tank drains through sand (stippled), through (E) an outflow that is raised to a level that would cause water to remain at the level of the dashed line, even if a siphon is interrupted to cause water to stop flowing into the main tank. (F) Plastic bottle and parts used to create small aquarium using the same logic as main aquarium. (G) Custom-built acrylic aquarium with drain in bottom and raised outflow with adjustable rate.

We also created a similar system for smaller experimental aquaria, but these appeared to be less effective long-term, as we saw high mortality in non-systematic studies. For the small aquaria (“condos”), we cut a large hole (∼5 cm diameter) into the bottom of 500 mL Nalgene™ Square PETG Media Bottles with Septum Closures (Figs. 2F, S7). We then modified a 100 μm cell strainer to fit in the bore of the media bottle in such a way that it would remain there when the closure of the media bottle was screwed back on, allowing water but not sand to flow through. We inverted the bottle and added a ∼3 cm layer of sand above the cell strainer, measured from the edge of the bottle cap. In plain terms, the condos are simply capped bottles, turned upside down, after creating a large hole in the bottom and a small hole in the cap. There is a strainer in the cap so water can flow down through the cap and not get clogged by sand. For these smaller aquaria, we used 5 mm diameter tubing as siphons from the reservoir tank and for the outflow. Similar to the large tank, the outflow tubes (placed in the septum closure) drained roughly halfway up the side of the aquaria to prevent the water from draining completely.

Most recently, we created custom aquariums from acrylic (Fig. 2G), with similar dimensions to the 5-gallon glass aquarium. We installed a bulkhead in the bottom and attached PVC tubing as a drain, which we again raised to about half the height of the aquarium to prevent complete drainage if inflow of water stops accidentally. We added a valve to control the rate of outflow and instead of a siphon, we added a float switch to set inflow rate. Currently, we raise the outflow pipe to set the level of water in the aquarium and use continuous inflow directly to the main aquarium, which works well as long as the water is draining freely. We use a flow rate of approximately 0.4 L/min.

We initially offered the ostracods multiple diet choices, including: (1) imitation crab, which is primarily pollock, (2) carnivore fish pellets or fish flakes, (3) tilapia fillets, and (4) anchovies, which have each made for successful food sources in Caribbean ostracods (Todd Oakley, personal observation; ^28, 43^). Although the adults appeared moderately interested in these food sources in non-systematic trials, we found the juvenile ostracods did not eat them, but they did actively feed on raw chicken liver, as do *Vargula hilgendorfii* individuals from Japan ^49^. In culture, we therefore provided pieces of chicken liver daily (∼1 cm^3^ per 200 animals). Due to the nocturnal life history of *V. tsujii*, we fed them in the late afternoon or early evening (around the beginning of the dark cycle), and removed the food from the aquarium in the morning (∼3-5 hours into the light cycle) to reduce the chance of bacterial fouling.

### Rearing of embryos

We tested two primary methods for rearing embryos. The first was less successful in terms of longevity of both the mothers and embryos, but provided usable data on embryogenesis. Regardless of rearing method, ostracods did not appear to eat while they were brooding.

For the first method, we kept females individually in circular wells in twelve-well dishes ^28^, with approximately 1 mL of 17^०^C seawater. We recorded visual observations of the females and their embryos and changed the water daily. The advantage of this method is ease of identification and observation of individual ostracods, allowing us to track the growth of specific embryos over a given period of time. However, all females which deposited embryos into the brooding chamber died before their embryos were released from the marsupium, although the precise cause of death is unknown. These embryos were then removed from the female marsupium by dissection, kept in wells, and observed until the animals hatched through the chorion (see Supplemental Video).

For the second method, we identified and removed brooding females from the aquarium and placed them in 50 mL Falcon™ tubes (2 females per tube). Roughly 5 mL of commercial aragonite sand for saltwater aquariums with most grain sizes 1.0-2.0 mm (Caribsea brand) was added to the bottom of each tube to allow females to burrow. Falcon tubes were then placed in a large seawater reservoir that ranged from 17-19°C. We changed the water in each tube daily, and imaged brooding females every other day with a camera attached to a dissecting scope.

We used key diagnostic features such as number of embryos and female size to distinguish individual females from one another. Of the thirteen individuals kept in Falcon^TM^ tubes, only one died prior to embryo release from the marsupium.

### Animal measurements and inferences of instar size classes

We imaged ostracods across all life stages using a standard dissecting microscope and an Olympus brand camera, and measured the carapace length, carapace height, eye width, and keel width with the software Fiji (Sample size = 626 individuals; Table S3) ^50^. These parameters are useful in the determination of the instar stages in other bioluminescent ostracods ^28^. A subset of these individuals (9 males, 11 females) were dissected to confirm sex. We partitioned the data into clusters using the KMeans clustering algorithm implemented in scikit-learn ^51^ in Python 2.7 ^52^. KMeans fits the data into k clusters, specified in the arguments, sorting each data point into the cluster which has the minimum euclidean distance to the centroid of the cluster.

**Table 3.** Additional measurements for further inferring A-I Male, A-I Female, and Adult Male instars, including length, height, eye width, and eye to keel distance (EKD) ± SD.

| Inferred Instar | N | Length | Height | Eye size | EKD |
| --- | --- | --- | --- | --- | --- |
| A-I Male | 15 | 1.566 ± 0.032 | 0.907 ± 0.039 | 0.218 ± 0.028 | 0.828 ± 0.053 |
| A-I Female | 15 | 1.592 ± 0.026 | 0.902 ± 0.030 | 0.198 ± 0.018 | 0.974 ± 0.038 |
| Adult Male | 13 | 1.647 ± 0.028 | 0.963 ± 0.023 | 0.242 ± 0.019 | 0.878 ± 0.040 |

The value for k was originally specified as eight based on the five known stages of instar development for related species of ostracods plus the embryonic stage and adult males and females. However, a k-value of eight clusters resulted in a split of the inferred adult female cluster into two clusters rather than the discrimination of adult males from inferred A-I instars. To avoid this apparent over-clustering, we ran our final analysis with a k-value of seven, which retained all inferred adult females in a single cluster but was unable to distinguish an adult male cluster from the inferred A-I instars. We assigned development stages to each cluster based on the relationship by comparison to the closely related *Photeros annecohenae*, which grew in 12-well dishes to allow more precise documentation of instars ^28^. We further split the A-I instar/Adult male cluster using the KMeans algorithm and a k-value of 3 with more precise measurements of length, height, and eye width, as well as eye to keel distance (EKD) from an additional sample of individuals that fell within this cluster (Sample size = 43). The three new clusters were then assigned to development stages by comparing eye size and shape parameters with what we expect based on *Photeros annecohenae* ^28^.

### Whole genome sequencing

We froze four adult female *Vargula tsujii*, collected from the WMSC Dock on 2017-09-01 in liquid nitrogen and shipped the specimens on dry-ice to the Hudson Alpha Institute for Biotechnology Genomic Services Lab (HAIB-GSL) for genomic DNA extraction and Chromium v2 Illumina sequencing library preparation. HAIB-GSL extracted the DNA using the Qiagen MagAttract Kit, yielding 141 ng, 99 ng, 153 ng, and 96 ng of extracted DNA respectively. This quantity of DNA was not sufficient for a size distribution analysis via agarose pulse field gel analysis, therefore we decided to proceed with Chromium v2 library prep with the 153 ng sample (sample # 5047), without confirmation of a suitable high-molecular weight DNA size distribution. We sequenced this library on three lanes of a HiSeqX instrument using a 151×151 paired-end sequencing mode, resulting in a total of 910,926,987 paired reads (260.6 Gbp). We attempted whole genome assembly of the resulting sequencing data with the Supernova v2.0.0 genome assembly software ^53^. We then used the Agilent Femto Pulse High Molecular Weight DNA capillary electrophoresis instrument at the MIT BioMicro Center to perform a post sequencing analysis of the size distribution of the DNA used for Chromium library preparation.

### Mitochondrial genome assembly

To achieve a full length mitochondrial genome (mtDNA) assembly of *V. tsujii*, we partitioned and assembled sequences separately from the nuclear genome. We mapped Illumina reads from our Chromium libraries (Table S3) to the known mtDNA of the closest available relative, *Vargula hilgendorfii* (NC_005306.1) ^54^ using bowtie2 (v2.3.3.1) ^55^ (parameters: --very-sensitive-local).

We then extracted read pairs with at least one read mapped from the resulting BAM file with samtools (parameters: view -G 12), then name-sorted, extracted in FASTQ format (parameters: samtools fastq) and input the reads into SPAdes (v3.11.1) ^56^ for assembly (parameters: --only-assembler -k55,127). Inspection of the resulting assembly graph using Bandage (v.0.8.1) ^57^, combined with a heuristic manual deletion of low coverage nodes (deletion of nodes with coverage <200x, mean coverage ∼250x) gave rise to two loops with sequence similarity to the *V. hilgendorfii* mtDNA that both circularized through a single path (Supplementary Figure S1). We then manually inspected this graph via a blastn (2.7.1+) ^58^ aligned against the *V. hilgendorfii* mtDNA through SequenceServer (v1.0.11) ^59^ and visualized with the Integrated Genomics Viewer (v2.4.5) ^60^. We noted that the single circularizing path was homologous to the two duplicated control regions reported in the *V. hilgendorfii* mtDNA ^54^. We surmised that this assembly graph structure indicated the *V. tsujii* mtDNA had the same global structure as the *V. hilgendorfii* mtDNA, and so used the Bandage “specify exact path” tool to traverse the SPAdes assembly graph and generate a FASTA file representing the circularized *V. tsujii* mtDNA with the putative duplicated control regions (Figure S1). To avoid splitting sequence features across the mtDNA circular sequence break, we circularly rotated this full assembly with the seqkit (v0.7.2) ^54, 61^ “restart” command, to set the first nucleotide of the FASTA file at the start codon of ND2. We then confirmed 100% of the nucleotides via bowtie2 re-mapping of reads and polishing with Pilon (v1.21) ^62^ (parameters: --fix all). We annotated the final assembly of the mitochondrial genome using the MITOS2 web server ^63^ with the invertebrate mitochondrial genetic code (translation table 5). We manually removed low confidence and duplicate gene predictions from the MITOS2 annotation.

## Results

### Population genetics analyses

Our COI haplotype network revealed minimal structure among *V. tsujii* populations in southern California (Figure 3A). Haplotypes from San Pedro and Catalina Island overlap or diverge by only one to three mutations. Haplotypes from San Diego individuals, although still very similar to haplotypes from Catalina Island and San Pedro, differ by at least eight mutations and show no overlap with haplotypes from other populations of *V. tsujii*. However, the haplotype network for 16S indicates that sequences from San Diego individuals are identical to some individuals from the other two populations (Figure 3B). Furthermore, although the model with three clusters was favored in the DAPC analysis for each gene (Supplementary Figures S2 and S3), membership in five of the six clusters includes individuals from different populations (Supplementary Figures S4 and S5). Therefore our analyses do not support three genetically distinct populations as defined by location.

**Figure 3.**
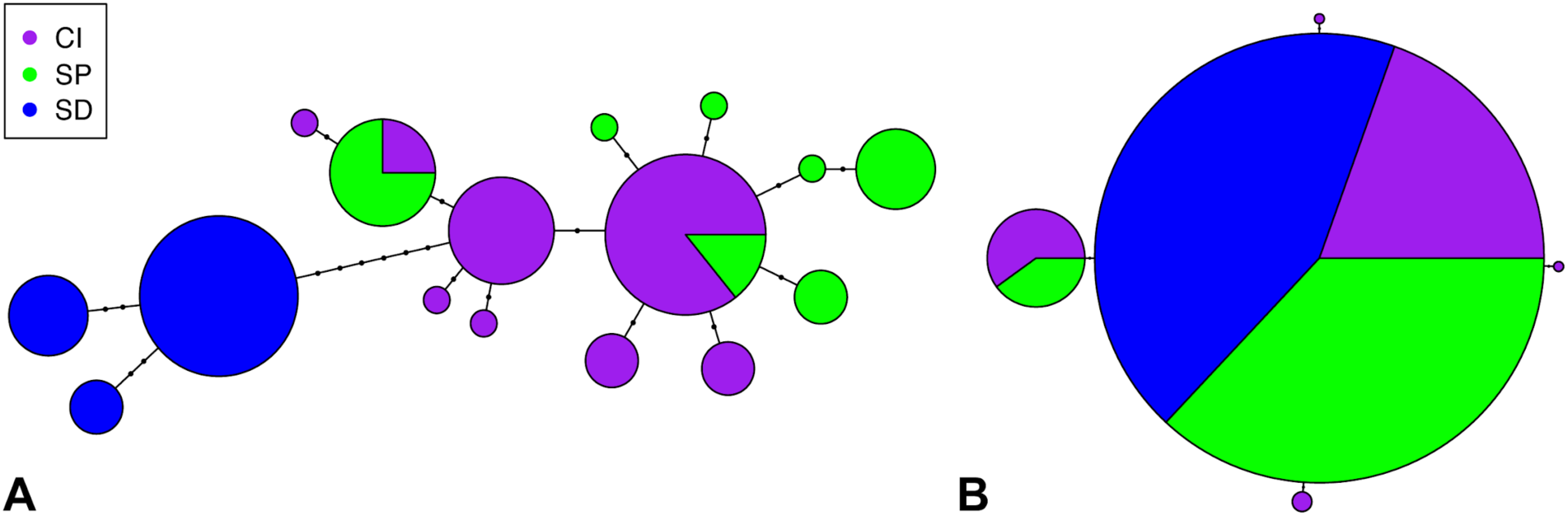
Haplotype networks for (A) COI sequences, and (B) 16S sequences of *Vargula tsujii* animals from three localities sampled. Abbreviations: CI, Catalina Island; SD, San Diego; SP, San Pedro.

### Embryogenesis Description

After fertilization and release into the marsupium, *V. tsujii* embryos resemble opaque, green, ovoidal embryos (Figures 4 and 5). After five to eight days of growth, separation of the yolk begins, which appears as a light green mass condensed within the embryo. Around ten to eighteen days after deposition into the brooding chamber, two very small red lateral eyespots are visible on each embryo and the separation of the yolk continues. Daily observations over the next six days showed continual growth and darkening of the eyes. By day sixteen the yolk is almost completely separated and has begun to form the gut of the ostracod. By days fifteen to nineteen, the eyespots are mostly black and the light organ (a light brown/yellow mass) appears anteroventral to the eyes of the embryo. Around twenty days after deposition, developing limbs in the form of two rows of small brown masses are visible along the ventral side of the embryos. At this point the gut has finished forming within the carapace and is visible as a yellow-white mass on the posterior end of each embryo. Limb formation continues over the next few days, and the eyes of each embryo become entirely black. Twenty-five to thirty-eight days following embryo deposition, embryogenesis is complete, and instars are released from the mother’s marsupium.

**Figure 4.**
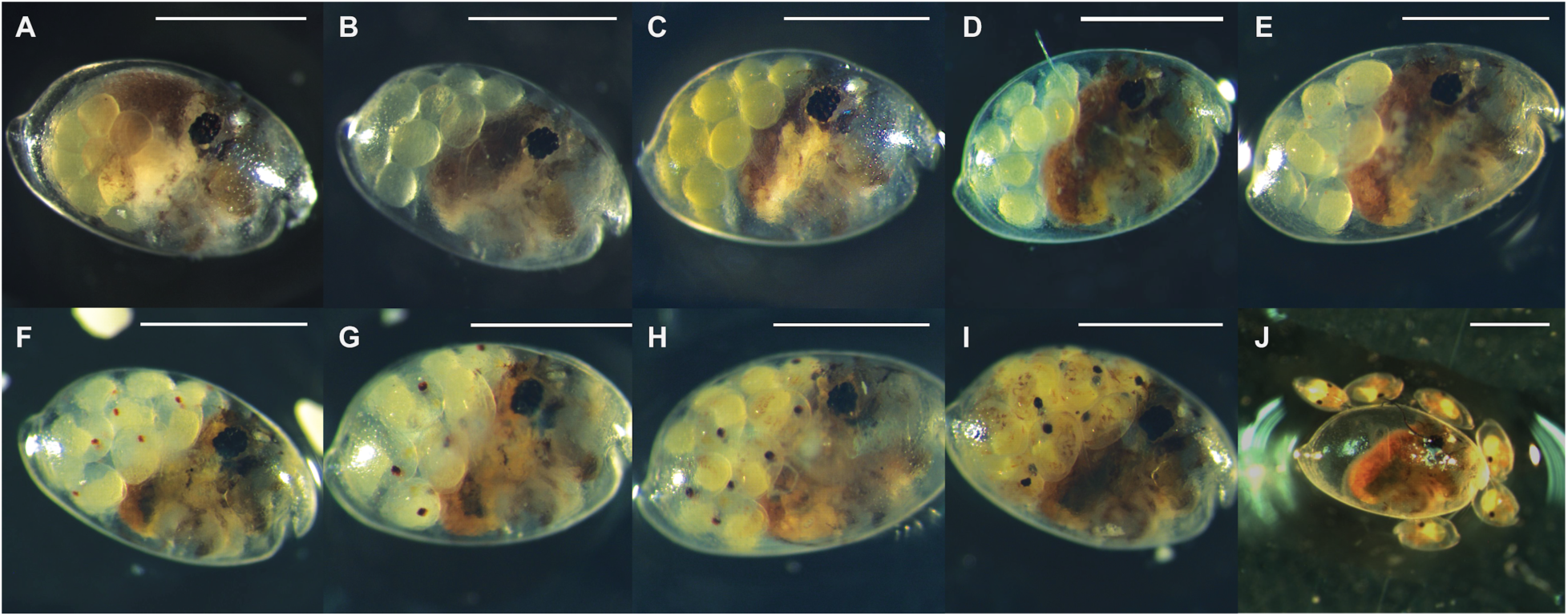
Embryogenesis within brooding females of *Vargula tsujii* post release into the marsupium. (A) Day 0, (B) Day 1-4, (C) Day 5-7, (D) Day 8-9, (E) Day 10-13, (F) Day 14-17, (G) Day 18-19, (H) Day 20-21, (I) Day 22-23, (J) Day 25-38.

**Figure 5.**
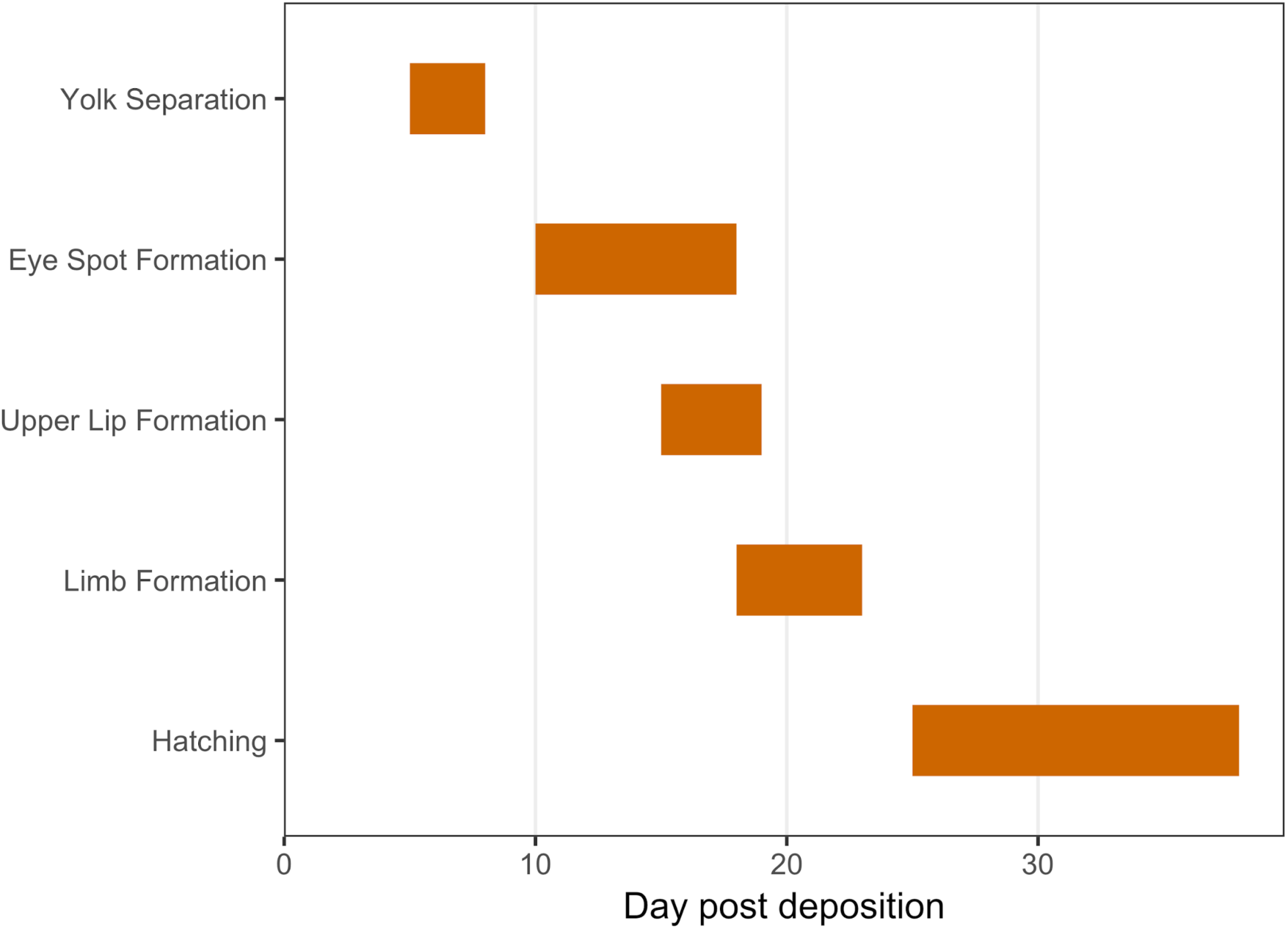
Summary of the timing of embryogenesis stages in *Vargula tsujii*. This represents the complete range of first appearance of each features across multiple broods, including: yolk separation (range = 5-8 days, N = 7); eye spot formation (range = 10-18 days, N = 12); upper lip formation (range = 15-19 days, N = 3); limb formation (range = 18-23 days, N = 11); hatching (range = 25-38 days, N = 11).

### Instar development

We inferred five juvenile instar stages (A-I, A-II, A-III, A-IV, A-V) and an adult stage for Vargula tsujii (Figures 6, 7A; Table 2). Laboratory measurements of specimens of unknown sex were compared with the measurements of sexed individuals (9 males, 11 females) to confirm interpretation of the instar stage of size classes (Figure 7C-D). Due to large amounts of overlap of A-I females with adult and A-I males, these three groups are currently indistinguishable in the large data set. However, when we attempted to distinguish these stages using additional measurements (particularly eye-to-keel distance, EKD), we find that EKD differentiates A-I males and females, and that length separates the A-I instars from adult males (Figures 6E-G, 8; Table 3). Sexual dimorphism in size becomes apparent in adulthood, with adult females clearly larger than adult males. However, compared to many other cypridinid ostracods, this dimorphism can be difficult to assess visually without a side-by-side comparison.

**Figure 6.**
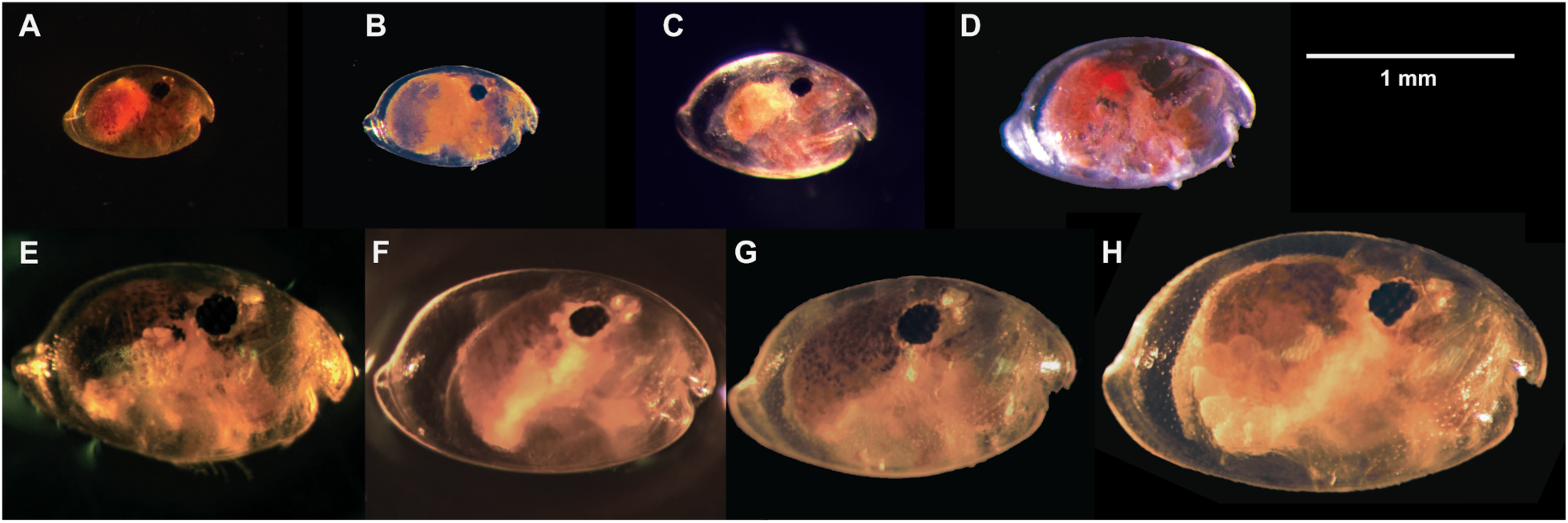
Images of selected individuals within each inferred instar stage: (A) A-V, (B) A-IV, (C) A-III, (D) A-II, (E) A-I male, (F) A-I female, (G) Adult male, (H) Adult female.

**Figure 7.**
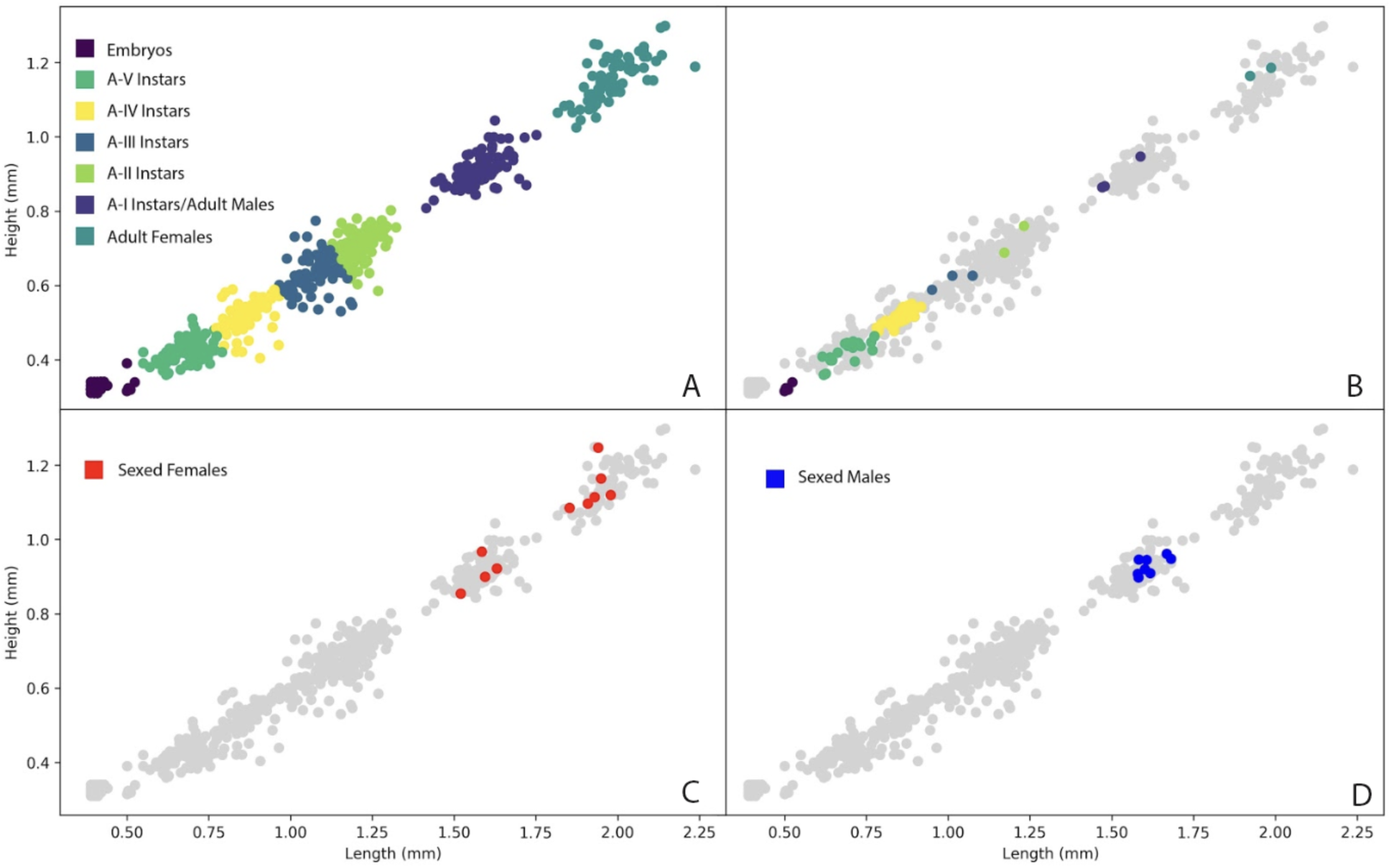
(A) KMeans clusters of *Vargula tsujii* developmental stages based on carapace height and width. Instar stages were assigned to each cluster via comparison to *P. annecohenae* and additional analysis of male and female size dimorphism. (B) Points representing laboratory cultured animals overlayed on wild caught and unknown data. Different colors represent different clusters in the k-means analysis, and are labeled with their inferred instar/adult stages. (C) Shows wild-caught animals dissected and identified as female in relation to the k-means clusters, and (D) shows wild-caught animals dissected and identified as male in relation to the k-means clusters.

### Evidence for complete life cycle in the lab

To assess whether a complete life cycle was obtained in the laboratory, we began with only brooding females in a small experimental aquarium. Those brooding females, although not tracked individually, released a number of juveniles, which we measured intermittently over time (Figure 6B). In this aquarium we found juveniles, and eventually adults, that fell across the full size range of wild-caught ostracods. After all of the broods hatched, we removed all adult females from the experimental aquarium to assess whether the ostracods would mature and mate in the laboratory. Following removal of the original adult females, we found adult brooding females that grew up in experimental aquarium that themselves gave birth to the next generation in 3-4 months time.

### Nuclear and mitochondrial genome assemblies

Our final *Vargula tsujii* nuclear genome assembly is highly fragmented, with an indication of high levels of heterozygosity between haplotypes (∼5%) and a high proportion of simple sequence repeats. Our post-sequencing analysis indicated the mode of the DNA size distribution for the four extracted DNA samples were 1244 bp, 3820 bp, 1285 bp (sample #5047), and 1279 bp, respectively, with a tail of ∼20-40 kbp in each sample.

The final *V. tsujii* mtDNA (15,729 bp) (Figure 9) aligned to the previously published *V. hilgendorfii* mtDNA with 98% coverage and 78.5% nucleotide identity, and is available on NCBI GenBank (MG767172). The *V. tsujii* mtDNA contains 22 tRNA genes, 13 protein coding genes, and 2 duplicated control regions in an identical positioning and orientation to the *V. hilgendorfii* mtDNA. The V. tsujii duplicate control regions (661 bp, 570 bp), are smaller than the homologous V. hilgendorfii control regions (855 bp, 778 bp). The alignment of the V. tsujii control regions to the V. hilgendorfii control regions indicate they possess reasonable nucleotide identity (∼73%), but the presence of unalignable regions ∼20% of the length of the regions suggests evolution constrained by size and/or GC% content rather than sequence identity alone. There is one polymorphic base in the final mitogenome assembly coding for a synonymous wobble mutation in the COII gene (COII:L76).

**Figure 9.**
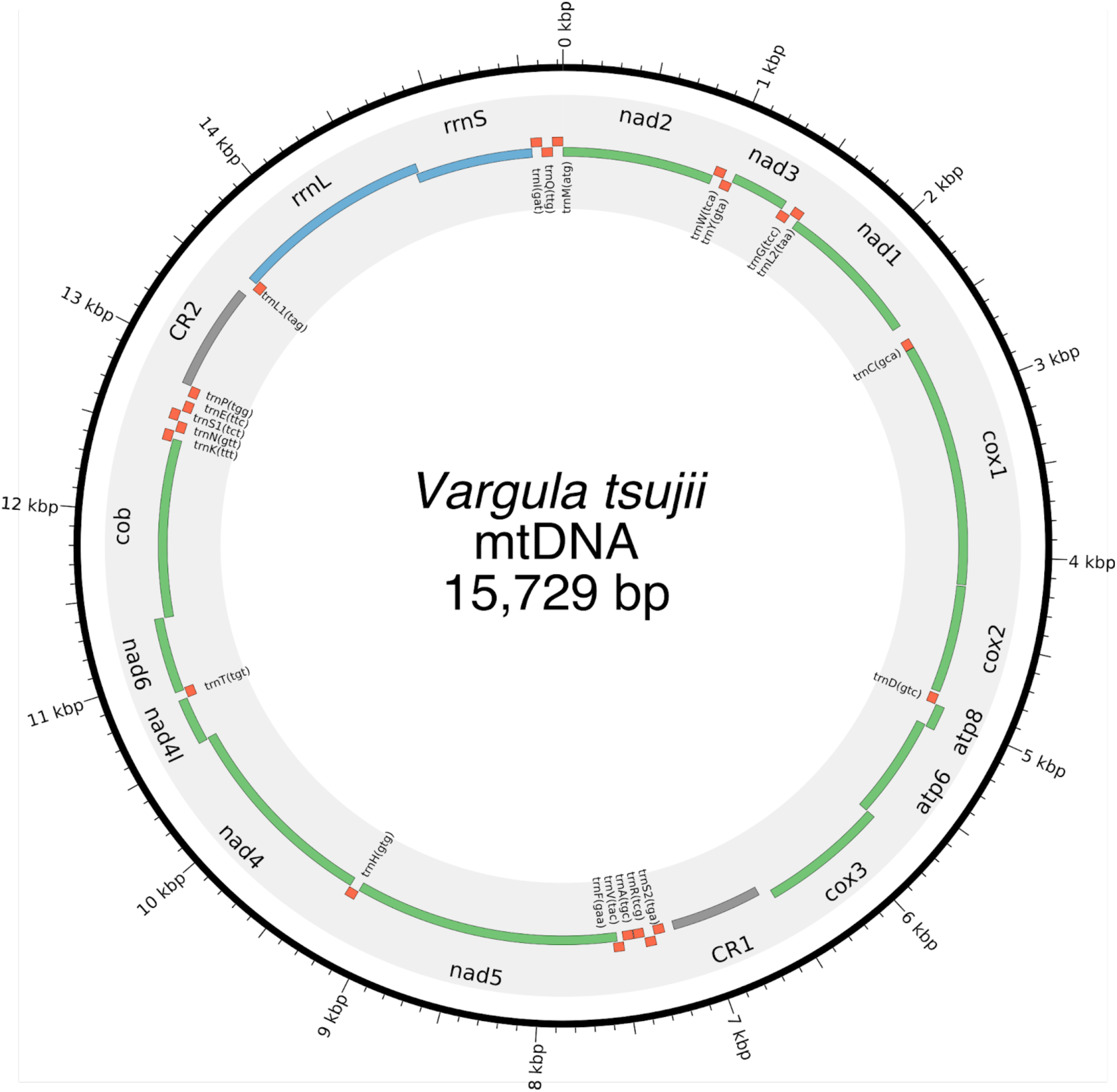
Mitochondrial genome of *Vargula tsujii*. Note the duplicated control regions, CR1 and CR2 (in gray). Figure produced with Circos ^78^.

## Discussion

Based on measurement data on the first and second generations of ostracods reared in the lab (Figure 5), and the presence of brooding female ostracods in our experimental aquarium where adult females had previously been removed, we can infer that we were successful in rearing *Vargula tsujii* through its complete life cycle in the lab. This is a first for any bioluminescent ostracod, as rearing in the lab has remained a significant hurdle for laboratory studies in these animals ^28, 29^. However, despite recent improvements, challenges still remain. Recent improvements include adding a neoprene gasket to the edge of the false bottom to prevent sand and detritus from falling below and fouling the aquarium and periodically siphoning the surface of the sand in the aquarium to reduce build-up of detritus and fouling microorganisms. But previous challenges mean our current population of animals is relatively small (between 500 and 1000 individuals), and we do not yet know if it is possible to continue the culture indefinitely without new input of animals. Currently, we input new wild-caught animals about every 6 months to increase the population size. Furthermore, the populations within condos were very prone to collapse, and animals do not survive long in 12-well dishes as they did in P. annecohenae ^28^, limiting current experimental work.

Our current population was cultured from animals collected near Wrigley Marine Science Center on Santa Catalina Island, which appears to contain much of the genetic diversity sampled across much of the known the range of *V. tsujii* (Figure 3). Our population genetics analyses support the hypothesis that populations of *Vargula tsujii* experience some gene flow across the localities sampled, including Catalina Island. These results are consistent with the hypothesis that *V. tsujii* represents a single, connected species. As such, we assume our culture of ostracods from Santa Catalina Island, and inferences made from it, may be representative of the genetic diversity of populations of *V. tsujii* across southern California coastal systems.

However, we cannot discount the possibility that there might be some variation in development times and instar sizes across the range of this species. Also of note is that this species is not easily collected throughout the reported range, and northern populations, if they exist, may be divergent from southern California populations.

### Attempted whole genome reconstruction

The 1C genome size of *Vargula tsujii* was previously reported to be 4.46 ± 0.08 pg (Female, n=3), and 4.05 pg (Male, n=1) ^64^, corresponding to a 1C genome size of 4.36 Gbp and 3.96 Gbp respectively and suggesting XO sex determination. Our reconstruction of the *Vargula tsujii* nuclear genome assembly is highly fragmented, with an indication of high levels of heterozygosity between haplotypes (∼5%) and a high proportion of simple sequence repeats. We speculate that our poor nuclear genome assembly results are likely due to the small size distribution of the input DNA, as larger fragment sizes are especially critical for the reconstruction of large genomes such as *V. tsujii*. Further protocol development is needed to extract suitable High Molecular Weight DNA, which could allow long-read sequencing that would improve assembly.

### The life cycle of Vargula tsujii

The embryogenesis of *Vargula tsujii* is similar to closely related species, and the 24-day average brooding duration falls within the range for other cypridinid ostracods (10-30 days) ^65^. The primary stages of *V. tsujii* late-stage embryogenesis are yolk separation, formation of eye spots, development of the upper lip, development of limbs, and emergence from the chorion and marsupium (Figure 5). In general, the late-stage embryonic development of *Vargula tsujii* is longer than that of other luminescent ostracods whose development has been described (Table 4). Longer overall development in the temperate V. tsujii is consistent with work in other crustaceans, which indicates that temperature has an inverse relationship with development time ^66^. However, earlier embryonic development in *V. tsujii* is faster than in the tropical *P. annecohenae*. In *P. anncohenae,* yolk separation begins on day 9 and eye development begins on days 14-15 ^28^, while those stages begin at 5-7 and 10-13 days, respectively, in *V. tsujii*. Additionally, the order of development in *V. tsujii* appears to differ from that of *Vargula hilgendorfii,* which starts to develop limbs on days 4-5, before eye and upper lip development ^67^. By contrast, *V. tsujii* limbs do not appear until after eye-spot and upper lip formation. The reasons for this difference are unclear.

**Table 4.**
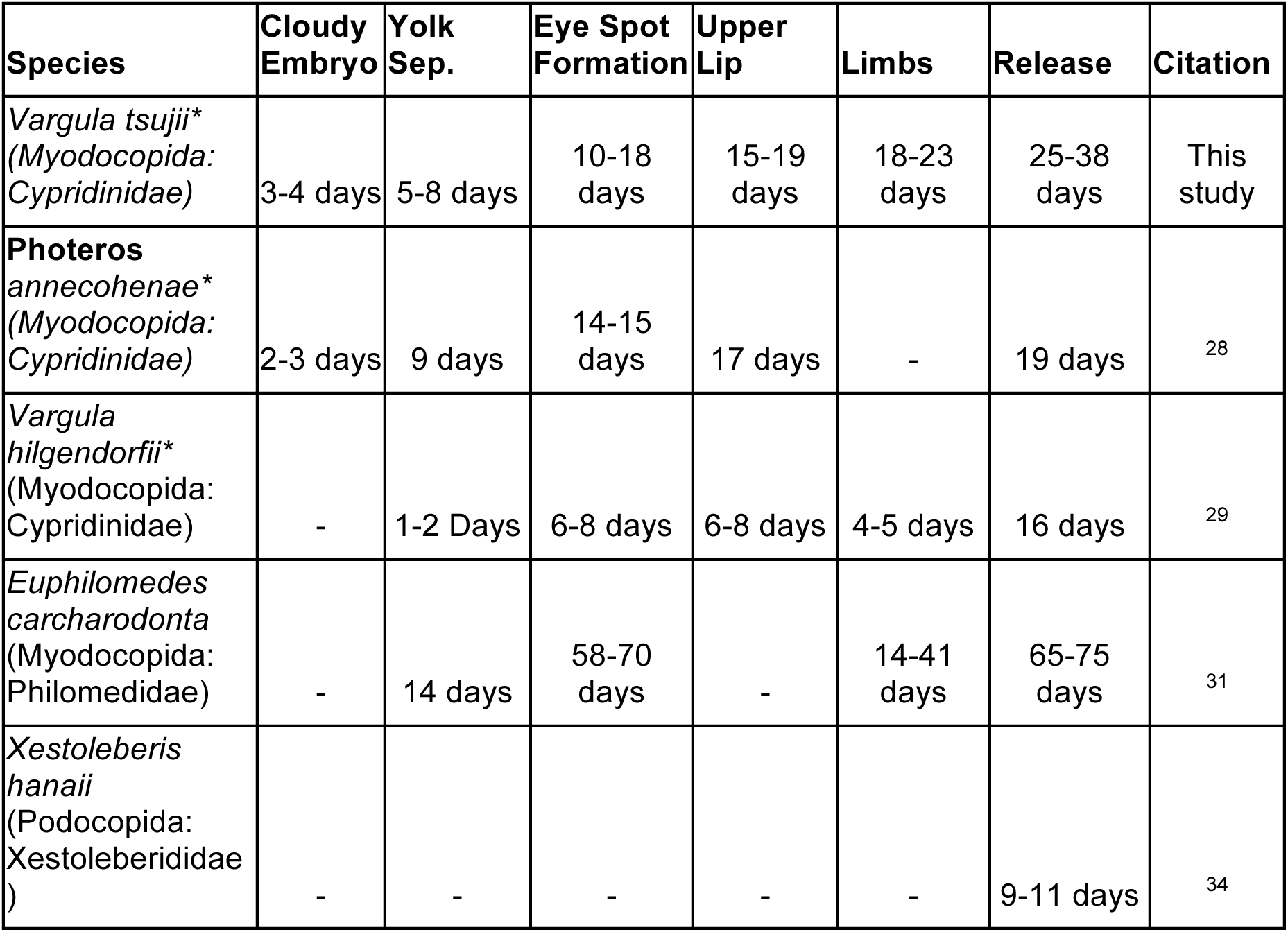
Comparison of embryogenesis timing between five different ostracod species post-deposition. Species with an asterisk (*) are bioluminescent cypridinids.

Juvenile development of *Vargula tsujii* is also quite similar overall to that of *Photeros annecohenae*, the only other bioluminescent cypridinid ostracod for which instar developmental data are available ^28^. We inferred five distinct instar stages (A-V through A-I), and A-I instars develop into size dimorphic adult males and females (Tables 2 and 3). It is important to note that these inferences represent hypotheses that can be tested by rearing and measuring individuals through molts, which we have not yet been able to accomplish in our culture. However, average measurements of size, including carapace length and height, as well as eye width, were comparable between respective instar stages in *V. tsujii* (inferred) and *P. annecohenae* (observed). However, in *V. tsujii* average eye width remains within the range of 0.060 mm to 0.0684 mm from the embryonic stage until the second instar, and makes a large jump by about two fold in the third instar (0.121 ± 0.0208 mm), as compared to a gradual increase in eye size in *P. annecohenae*. In our KMeans clustering analysis, the second-to-largest cluster of *V. tsujii* appears to contain adult males as well as A-I males and females. This finding differs from that in *P. annecohenae*, where adult males were found to cluster separately from A-I males and females and adult females based on size and shape, and A-I males and females could be distinguished by certain shape parameters, as well as eye size ^28^. In V. tsujii, A-I males and females appear to be differentiated largely by eye-to-keel distance, which was not measured in previous studies ^28^, and adult males can be separated from A-I instars by length (Figure 8; Table 3). We hypothesize that this apparent reduction of sexual dimorphism observed in the later stages of development, compared to *P. annecohenae*, may occur because *V. tsujii* may have lost the ability to use luminescence for courtship ^23^, so selection acting on differences in male and female shape may have been reduced.

**Figure 8.**
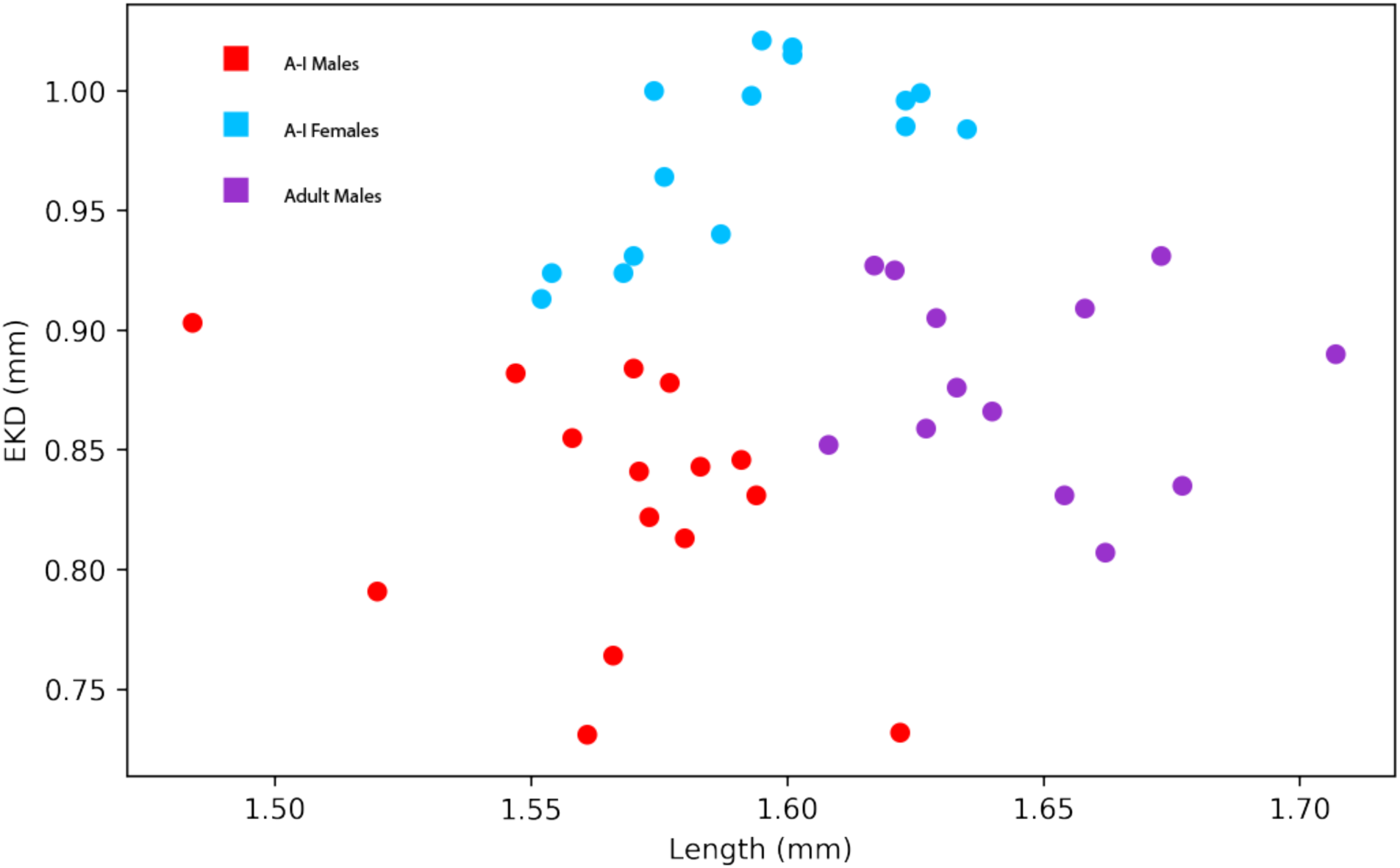
Plot of length versus eye to keel distance (EKD) in animals assigned to the A-I Instar/Adult Male cluster in the previous KMeans analysis (Figure 7A). These points were then clustered via KMeans analysis into three clusters, which we assigned to A-I Males (red), A-I Females (blue), and Adult Males (yellow).

### Benefits of completing the life cycle

Cypridinid ostracods represent an independent origin of endogenous autogenic bioluminescence. This group is a natural comparison to other, well-studied bioluminescent courtship systems, such as that of fireflies ^68^. Other possible laboratory models for bioluminescence, such as brittle stars ^69^ and cnidarians ^70^ are useful for some research tasks, such as studies of luciferins obtained from the diet (e.g., coelenterazine). Others might be easier to rear in the laboratory, such as the self-fertilizing brittle star *Amphiolus squamata* ^71^. However, ostracods produce their own luciferase and luciferin, unlike brittle stars and cnidarians.

Ostracods also have relatively fast generation times (on the order of months), which can be modified somewhat by adjusting the temperature of the water, as is common among ostracods ^27–29, 72, 73^. In comparison, fireflies are usually on a six-month to one-year life cycle that is difficult to alter ^74–76^, and some brittle stars appear to spawn only seasonally ^77, 78^. Furthermore, culture of an organism (*Vargula tsujii*) from a clade with an independent origin of autogenic bioluminescence allows for comparative studies between luminescence in ostracods and the autogenic bioluminescence found in fireflies ^16^ or other organisms, which would not otherwise be possible.

Ostracods will be incredibly useful for studying the physiology and biochemistry of bioluminescence ^2, 16^, as both diversification and behavioral changes in cypridinids related to luminescence have occurred using the same luciferin and homologous luciferases ^25^. Furthermore, although Vargula tsujii itself does not possess bioluminescent signaling, we expect a similar culturing system could be useful for some signaling species as well, at least those that come to baited traps. Given that ostracod courtship signals may be simulated in a laboratory setting, this system could allow us to assess male and female behavior in response to changes in signal parameters, such as intensity, color, and pulse duration ^79^. These tools will provide researchers with an excellent system for asking important questions related to selection on genes important for rapid diversification ^80^, including those related to behavior ^81^, sexual selection ^82^, and the role of the biochemical properties of bioluminescence ^26^.

Importantly, the biosynthetic pathway for luciferin production is currently unknown for any animal. One major obstacle to describing this pathway has been building laboratory cultures for animal species that produce luciferin themselves and also have a relatively fast generation time. Solving this bottleneck to accessing specimens for bioluminescent ostracods has set the stage for investigations into the biosynthetic pathway of cypridinid luciferin and its possible use in biomedical tools such as optogenetics ^83^. Finally, this is an excellent system with which to consider developing genome editing techniques (e.g., CRISPR/Cas9) ^84^ to test candidate genes for luciferin biosynthesis and to assess how genetic variants in critical genes might affect production or perception of bioluminescent signals.

## Acknowledgements

We thank the USC *Wrigley Institute* for *Environmental Studies* for transportation and use of the facilities at the Wrigley Marine Science Center on Catalina Island, CA. JDM worked on the project at Wrigley during the Catalina Marine Biology Semester hosted by California State University and with support from the NSF (Award DEB·1457462 to ET). JAG was supported by the NSF (PRFB Award: 1711201). Population genetic work was supported by a mini-grant from Cal State LA to ET; sequence data were obtained from the Masters thesis of KH Lee; DM Jacinto and E Stiner helped with collections throughout California; RM Serrano helped with measurements of some specimens; N Hensley contributed data on collecting locations. Genomic sequencing was supported by the Beckman Young Investigator Award to JKW from the Arnold and Mabel Beckman Foundation and by a UCSB Faculty Seed Grant to THO. Finally, we would like to thank Simone Brandão and two anonymous reviewers for their comments on our manuscript.

## Author contributions

JAG, ET, JKW, and THO conceived of and designed the study; JAG, GM, MNB, MSD, JDM, TRF, and DTS contributed to the acquisition and analysis of the data in this study; all authors participated in the interpretation of data, helped draft or substantially revise the manuscript, and gave final approval for publication.

## Competing interests

The authors declare no competing interests.

## Data availability

Sequence data for COI and 16S are available in GenBank (Table S1) and our whole genome Chromium reads are available in the Sequence Read Archive (SRR9308458). Scripts and data for the population genetics and k-means clustering analyses are provided on Github (https://github.com/goodgodric28/vargula_tsujii_culture). An additional video which shows *Vargula tsujii* hatching from the chorion and outside the mother is provided on Youtube (https://youtu.be/8o5CqHTcTjI).

## Supplementary Materials

**Table S1.** List of collections locations or attempted collection locations for *Vargula tsujii*. Collection locations for which a latitude and longitude could not be determined were not figured, but are included here.

**Table S2.** List of specimens examined in this paper for which sequences were obtained, including locality, museum collection numbers (where applicable), and GenBank accession numbers for the two genes sequenced (COI and 16S).

**Table S3.** Raw measurements of *Vargula tsujii* individuals, including carapace height and length, keel width and eye width.

**Table S4.** Unfiltered library sequencing statistics for *V. tsujii*.

**Figure S1.** Traversal of the cleaned SPAdes *V. tsujii* mitochondrial genome assembly graph nodes plotted by Bandage (gray). The single circularizing path representing the collapsed duplicate control regions are shown in blue. The black stroke shows the path traversed through the assembly graph followed to produce the circularized mtDNA FASTA file with two control region, matching the structure of the V. hilgendorfii mtDNA (Ogoh and Ohmiya 2004).

**Figure S2.** Results for our 16S DAPC model test to determine the number of clusters that is favored via comparison of Bayesian Information Criterion (BIC) values.

**Figure S3.** Results for our COI DAPC model test to determine the number of clusters that is favored via comparison of Bayesian Information Criterion (BIC) values.

**Figure S4.** Cluster membership based on our 16S DAPC analysis compared with the original population designation for each specimen. Overall, the clusters are not representative of any population-level structure.

**Figure S5.** Cluster membership based on our 16S DAPC analysis compared with the original population designation for each specimen. Overall, the clusters are not representative of any population-level structure.

**Figure S6.** Exploded diagram of custom aquarium for rearing *Vargula tsujii*. A photo of built aquarium is in main text, Figure 2. **A**. Plumbing components include 1. PVC pipe 1.5 inch diameter. 2. PVC socket ball valve to control the rate of outflow 3. PVC 1.5 inch elbows 4. PVC elbow with one side threaded to screw into 5. threaded bulkhead to span bottom of aquarium for drainage 6. float switch valve to control water level (not shown on Figure 2 of main text). **B**. Acrylic panels of ¼ inch thickness welded together with solvent include 7. Two panels of dimension 16.5 x 13.25 inches that form the long sides of the aquarium 8. Two panels of dimension 8 x 10.75 inches that form the short sides of the aquarium. One of these panels has a hole drilled for the water inflow and float switch 9. One panel of 16.5 x 7.5 inches to form the bottom of the aquarium, including a hole drilled for the bulkhead 10. One panel of 16.5 x 7.5 inches to forms a false bottom for the aquarium similar to the bottom of an undergravel aquarium filter. We drilled small holes throughout to allow water but not sand to flow through. (Not shown on Figure 2). In practice, we welded legs on the false bottom to allow it to stand on the true bottom of the aquarium above the height of the bulkhead drain.

**Figure S7.** Small aquarium (“condo”) setup using an inverted 500 ml bottle. a. Seawater feed through a hose into reservoir tank. b. Inverted bottle with hole in bottom and tube piercing septum cap c. Siphon to bring water to condo (note in this staged example for photo, reservoir tank level is higher than condo, so water would flow out the top of the condo) d. Reservoir tank receives water from inflow and overflows e. Stand to hold condos f. Outflow tube using ¼ inch aquarium tubing, raised to set water level.

## Notes

https://github.com/goodgodric28/vargula_tsujii_culture

https://youtu.be/8o5CqHTcTjI

